# Microbiota-dependent histone butyrylation in the mammalian intestine

**DOI:** 10.1101/2022.09.29.510184

**Authors:** Leah A. Gates, Bernardo Sgarbi Reis, Peder J. Lund, Matthew R. Paul, Marylene Leboeuf, Zara Nadeem, Tom S. Carroll, Benjamin A. Garcia, Daniel Mucida, C. David Allis

## Abstract

Posttranslational modifications (PTMs) on histone proteins are a key source of regulation on chromatin through impacting genome organization and important cellular processes, including gene expression. These PTMs often arise from small metabolites and are thus impacted by cellular metabolism and environmental cues. One such class of metabolically regulated PTMs are histone acylations, which include histone acetylation, along with butyrylation, crotonylation, and propionylation. We asked whether histone acylations of intestinal epithelial cells (IECs) are regulated through the availability of short chain fatty acids (SCFAs), which are generated by the commensal microbiota in the intestinal lumen. We identified IECs from the cecum and distal mouse intestine as sites of high levels of histone acylations, including histone butyrylation and propionylation. We identified specific sites of butyrylation and propionylation on lysine 9 and 27 on histone H3. We demonstrate that these acylations are regulated by the microbiota, whereas histone butyrylation is additionally regulated by the metabolite tributyrin. Furthermore, we identify tributyrin-regulated gene programs that correlate with histone butyrylation and demonstrate that histone butyrylation (H3K27bu) is associated with active gene regulatory elements and levels of gene expression. Together, our observations demonstrate a physiological setting in which previously uncharacterized histone acylations are dynamically regulated and associated with gene expression.

## INTRODUCTION

Cells are exposed to a variety of stimuli in the environment, which can induce signals that converge onto chromatin and impact subsequent cellular processes. Posttranslational modifications (PTMs) on histone proteins, or histone marks, represent one major way in which the chromatin landscape is dynamically regulated. Some of these PTMs have crucial regulatory roles in processes such as transcription, chromatin organization, and more (Allis & Jenuwein, 2016). Understanding how these PTMs are regulated and their functional roles are thus critical for determining molecular mechanisms of fundamental biological processes such as gene regulation.

Recently, different types of chemical modifications have been identified on histones that greatly expand the diversity of histone PTMs (Farrelly et al., 2019; Tan et al., 2011; D. Zhang et al., 2019). However, the physiological functions of many of these PTMs have not yet been defined. Here, we have focused our studies on histone acylations, which include histone acetylation, butyrylation, crotonylation, and propionylation (Sabari et al., 2016). Histone acylations are structurally closely related PTMs that are generally associated with gene expression and their levels can be regulated through cellular metabolism (Brownell et al., 1997; Goudarzi et al., 2016; Kebede et al., 2017; Sabari et al., 2015, 2016). Furthermore, histone acetylation has been linked to transcriptional dynamics in the gut and liver in the context of circadian rhythm changes (Tognini et al., 2017). We thus wished to further investigate potential physiological roles of these regulatory PTMs *in vivo*.

The donor molecules of the majority of histone PTMs are small metabolites. Histone PTMs are thus not only regulated by enzymes that deposit or remove this modification, but also by the availability of donor molecules in the nucleus (Carey et al., 2015; Dai et al., 2020; Wellen et al., 2009). Short chain fatty acids (SCFAs) are metabolites that are generated at high concentrations in the intestinal lumen, through the action of microbes that ferment fiber (Koh et al., 2016). Specifically, acetate, butyrate, and propionate are estimated to accumulate in the human intestinal lumen to millimolar concentrations (Cummings et al., 1987). These metabolites can then act on neighboring intestinal epithelial cells (IECs) or distal tissues via circulation. Many studies have investigated how SCFAs bind to receptors, feed into energy pathways, and can act as inhibitors of histone deacetylase (HDAC) enzymes (den Besten et al., 2013; Brown et al., 2003; Donohoe, Collins, et al., 2012; Fleming et al., 1991; Poul et al., 2003; Roediger, 1982; Vidali et al., 1978). In addition, acetylation can be traced from bacterial fermentation of fiber to be deposited onto histones as PTMs (Lund et al., 2021). Alterations in the commensal microbiota or SCFA levels have also been demonstrated to regulate select histone acetylation and methylation PTMs, as well as crotonylation (Fellows et al., 2018; Krautkramer et al., 2016). Together, these studies suggest that metabolites can regulate the chromatin landscape and that there is precedent for the microbiota impacting select histone PTMs.

In this study, we aimed to determine physiological roles of histone acylations, predominantly focusing on the less characterized PTM of butyrylation. We hypothesized that since the gut harbors high concentrations of butyrate and other SCFAs, this may be a particular site where these metabolites may be deposited *onto* chromatin and have functional roles as histone acylations in gene regulation in the intestine.

## RESULTS

In order to investigate the role of histone acylations, especially non-acetyl acylations, in a physiological setting, we decided to first identify where these modifications occur throughout different tissues in the body. We hypothesized that tissues may harbor different levels of histone acylations depending on their local environment, and that the gut in particular may be an environment conducive to histone acylations due to the levels of SCFAs (Figure 1A). We thus purified histones from abdominal tissues from mice using acid extraction. We primarily focused on different regions of the gut in our tissue array, and then probed these histones for different PTMs (Figure 1B). We used pan-lysine modification antibodies, which we reasoned would recognize a variety of lysines modified with our PTMs of interest independent of the specific positions of the modification sites. Interestingly, we found that the cecum and large intestine displayed higher levels of histone butyrylation and propionylation compared to other tissues. These findings are in accordance with the fact that the cecum and colon are the major sites of microbial fermentation of dietary fiber and production of SCFAs (den Besten et al., 2013). We therefore decided to focus our studies on the cecum and colon for subsequent experiments.

**Figure 1:**
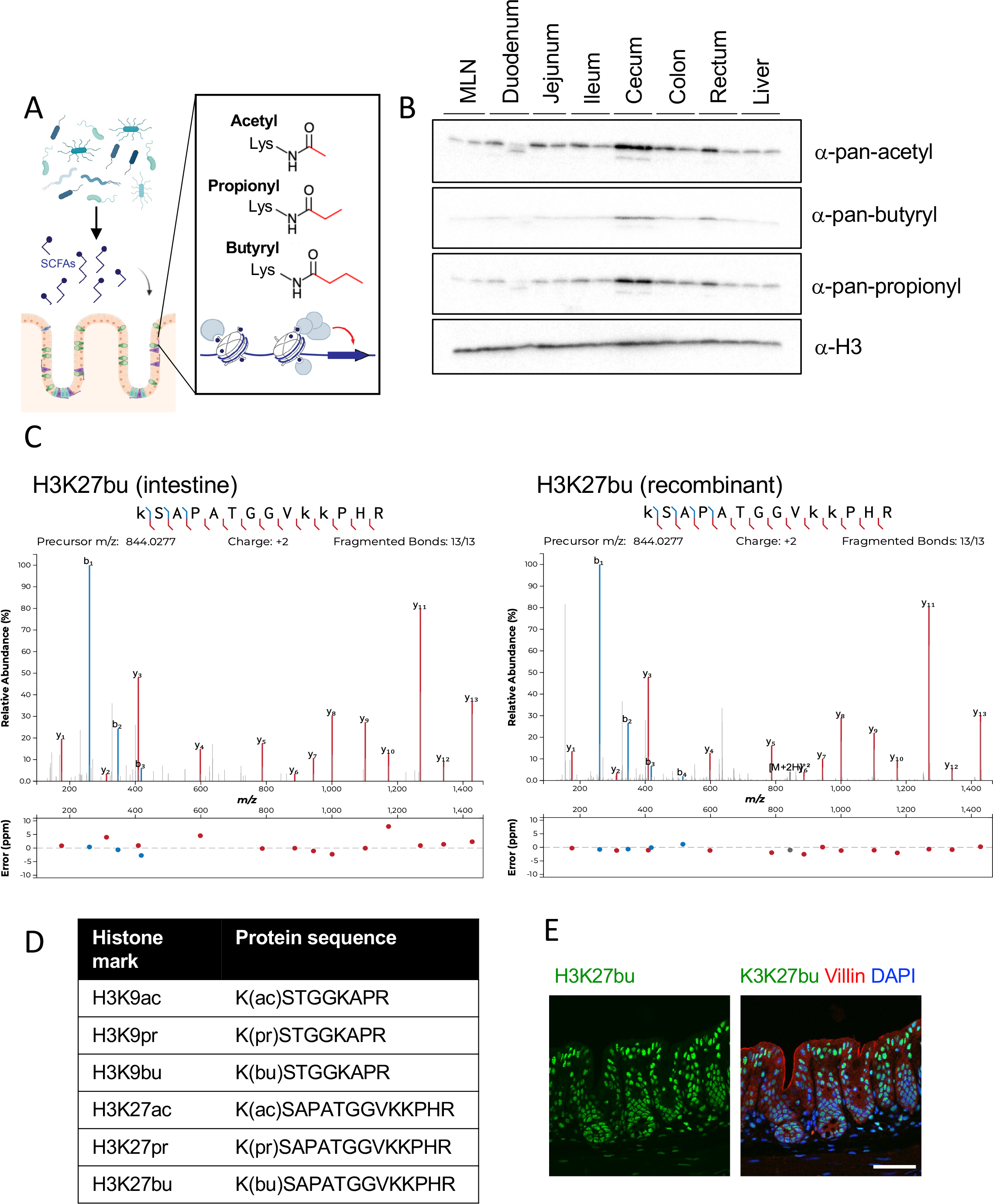
The intestine is an environment that harbors a variety of histone acyl marks. (A) Schematic of overall hypothesis and the structure of select histone acylations. (B) Tissue distribution of histone acylations. Histones were purified from tissues from C57BL/6 mice (n = 2 shown) using acid extraction and then subject to immunoblotting with pan-lysine antibodies targeting different acylation modifications. H3 serves as a loading control. (C) Characterization of specific sites of histone acylation. LC-MS/MS analysis of extracted histone from the mouse cecum. Representative fragmentation spectra are shown for endogenous H3K27bu in the intestine (left) and for recombinant H3K27bu on modified nucleosomes (right). (D) Table summarizing the different histone acyl marks detected in the mouse intestine. (E) H3K27bu displays bright staining in the intestinal epithelium. Representative immunofluorescence images of H3K27bu, Villin (marker of epithelium), and DAPI in mouse cecal sections, n = 5 C57BL/6 mice. Scale bar = 50 μm.

We next aimed to identify specific sites of histone butyrylation and propionylation in the mouse gut. To this end, we performed mass spectrometry on the extracted histones from the mouse cecum. We identified four specific PTMs with high confidence: butyrylation and propionylation on histone H3 lysine 27 (H3K27br and H3K27pr, respectively) and histone H3 lysine 9 (H3K9bu and H3K9pr) (Figure 1C,1D, S1A). Previous studies detected crotonylation in the colon (Fellows et al., 2018), however we did not observe crotonylation in the cecum by mass spectrometry. We noticed during our mass spectrometry analysis that the acylations on H3K27 displayed stronger signals than those on H3K9, which potentially could be due to differences in abundance and/or ionization properties. We also utilized recombinant nucleosomes to examine the dose-response behavior of the different histone acyl marks when analyzed by mass spectrometry (Figure 1C, S1C). Upon testing a dilution series of acylated nucleosomes spiked into unmodified nucleosomes, we observed that H3K27bu produces a lower signal than H3K27ac at similar relative abundances and that the butyryl mark for both H3K9 and H3K27 becomes undetectable at levels where the acetyl mark remains visible (Figure S1C). This suggests that the abundance of histone butyrylation and perhaps other acylation marks may be underestimated unless synthetic standards are utilized for quantification.

Since acyl modifications are structurally closely related, we next tested antibodies targeting these PTMs for selectivity (Supplemental Figure 2). We first utilized recombinant nucleosomes to test butyryl and acetyl antibodies, which displayed selectivity towards their specific modifications. The antibody targeting H3K27bu shows some cross-reactivity with H3K9bu and crotonylation, but minimal cross-reactivity with H3K27ac (Figure S2A). Since our mass spectrometry results did not detect histone crotonylation (data not shown), we concluded that this was satisfactory for our purposes of investigating histone butyrylation in the intestinal tract. To test histone propionylation antibodies, we utilized dot blotting with peptides, since commercial propionylated nucleosomes were not available (Figure S2B). The antibody targeting H3K9pr appeared selective, with minimal cross-reactivity towards H3K9bu and H3K9ac. The H3K27pr antibody was the least selective, as it had almost equal intensity to recognize H3K27bu and H3K27pr, and had minimal cross-reactivity with H3K9bu. Due to both the antibody specificity and signal intensity of H3K27bu, we decided to focus our efforts on this particular modification. To further characterize the localization of H3K27bu in the mouse intestine, we stained intestinal sections from the mouse gut to visualize this histone mark in the setting of the complex tissue. We found that H3K27bu is widespread throughout the cecum sections and displays bright staining in the intestinal epithelia (Figure 1E). These data suggest that these select non-canonical histone acylations, specifically H3K27bu, are in intestinal epithelial cells in regions that harbor fermenting bacteria.

**Figure 2:**
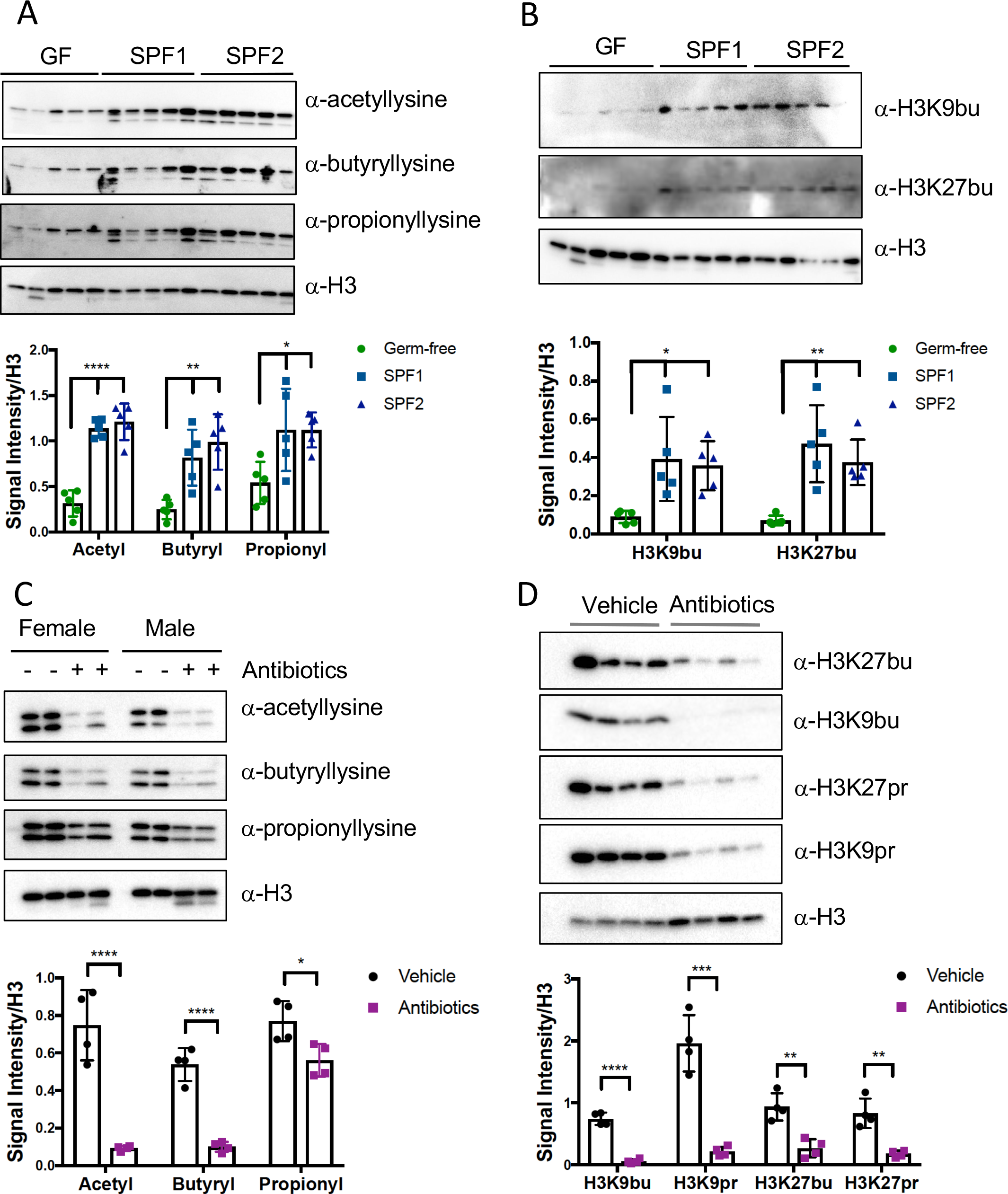
Histone acyl marks, including select non-acetyl acyl marks, are dependent on the microbiota. Immunoblotting (top) and quantification of signal intensity relative to histone H3 (bottom). Histones were extracted from the cecum and each lane represents an individual mouse. Quantification was performed in ImageJ and H3 serves as a loading control. (A) Pan-acylations and (B) specific histone acylations are reduced in germ-free mice. Germ-free (GF) mice (n = 5), conventional mice (SPF = specific pathogen free, SPF1 = C57BL/6NTac, n = 5, SPF2 = C57BL/6J, n =5). * p < 0.05, ** p < 0.01, **** p < 0.0001 by one-way ANOVA and adjusted for multiple comparisons. (C) Pan-acylations and (D) specific histone acylations are reduced in antibiotic-treated mice. C57BL/6J mice were treated with (n = 4) or without (n = 4) antibiotics (ampicillin, vancomycin, neomycin, and metronidazole) for seven days. * p < 0.05, ** p < 0.01, *** p < 0.001, **** p < 0.0001 by Student’s t-test.

Since we observed high levels of histone acylations in the mouse gut, we next investigated whether these PTMs were regulated by the commensal microbiota. We first extracted histones from germ-free or conventional mice and found that germ-free mice had reduced levels of histone acetylation, butyrylation, and propionylation (Figure 2A-B). We also sought to investigate whether this same type of regulation of the chromatin landscape would occur in mice that have an already established commensal microbiota. Therefore, we performed short-term treatment of conventional mice with an antibiotic cocktail and observed reduced levels of these same PTMs (Figure 2C-D). Together, these data suggest these histone acylations are sensitive to changes in the commensal microbiota *in vivo*.

While we demonstrated that alterations in the microbiota impact these histone PTMs, we wondered whether the microbial metabolites themselves were sufficient to regulate histone acylations. To answer this question, we set up a system in which we could reduce the microbiota and add an exogenous metabolite to the mouse gut (Figure 3A). We treated mice with or without ampicillin, which as a sole agent is sufficient to reduce select histone acylations. Then, we orally gavaged mice with an analog of butyrate, tributyrin, or vehicle control. We chose to use tributyrin instead of sodium butyrate, since it has been reported to be absorbed in the cecum and colon of mice versus the duodenum (Kelly et al., 2015). Importantly, tributyrin treatment was able to partially rescue histone butyrylation in the cecum (Figure 3B), suggesting that a specific metabolite alone is at least partially sufficient to regulate these PTMs in the intestine.

**Figure 3:**
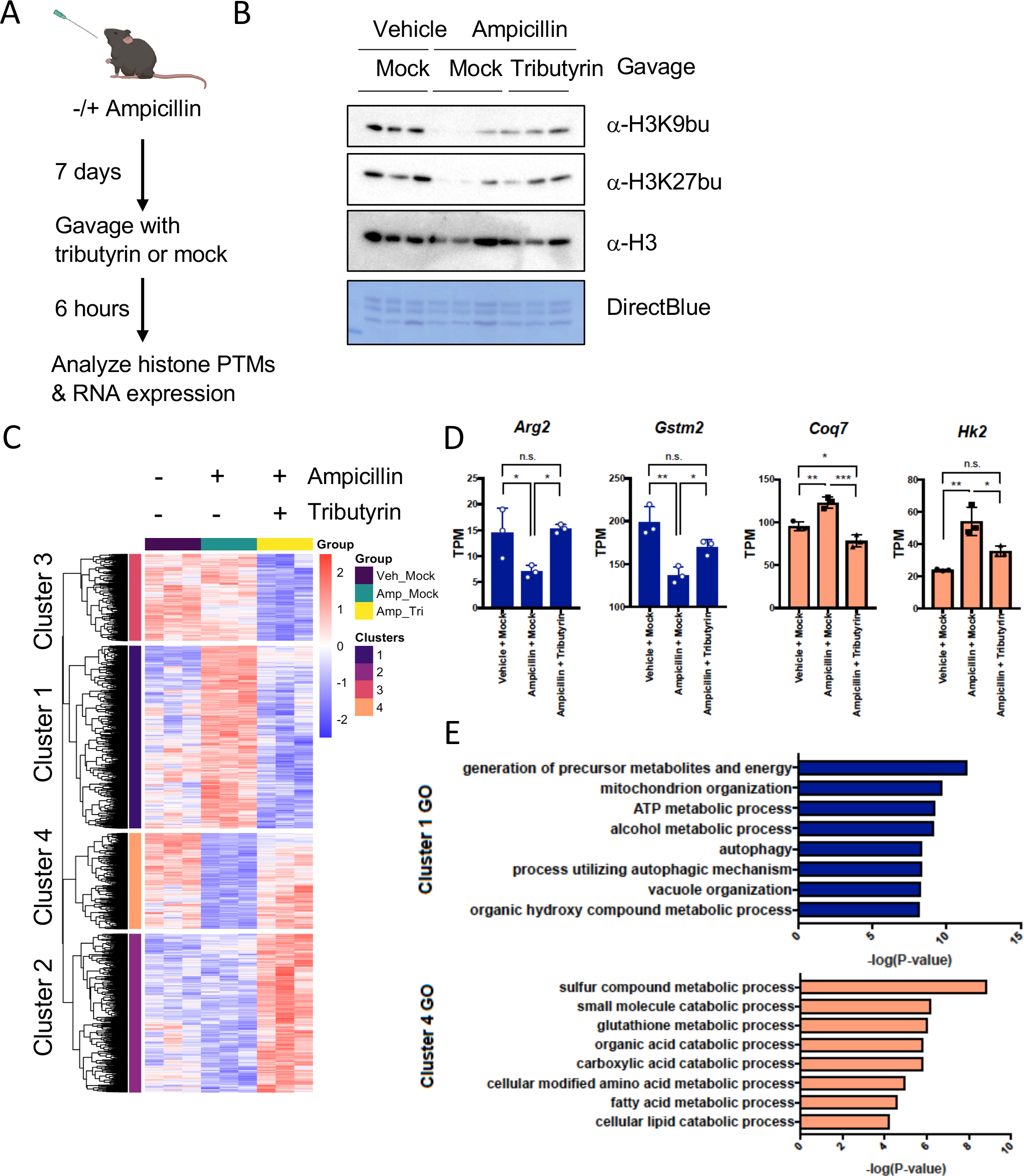
Microbial metabolites regulate select histone acyl marks and gene expression. (A) Schematic of experimental system. Mice are treated with or without Ampicillin for seven days, followed by gavage with vehicle or tributyrin. (B) Histone butyrylation dynamically changes with butyrate treatment. Western blot showing levels of histone butyrylation with different treatments. Representative experiment shown, n = 3 per group. (C) Heatmap of significant genes changing with tributyrin treatment by RNA-seq. Significant genes changing between mice treated with ampicillin plus tributyrin vs. ampicillin with mock gavage, padj < 0.05 by pairwise test, are shown. Hierarchical clustering was performed after determining optimal number of clusters using the elbow method. The heatmap intensity corresponds to the rlog counts as shown in the legend. Individual mice serve as biological replicates, n = 3 per group. (D) Visualization of expression of select genes in clusters 1 (blue) and 4 (red). Error bars = SEM. * p-value = < 0.05, ** p-value = < 0.01, *** p-value = < 0.001 by one-way ANOVA and adjusted for multiple comparisons. (E) Gene ontology (GO) of significant differentially expressed genes in clusters 1 and 4. Top eight most significantly enriched categories are shown.

We then analyzed gene expression changes in intestinal epithelial cells (IECs) when mice are treated with antibiotics and tributyrin, in order to address potential functional consequences of altering histone butyrylation. After treating mice and isolating IECs, we isolated RNA and performed sequencing. Our RNA-seq results indicated that ampicillin and ampicillin plus tributyrin induced robust changes in gene expression in IECs (Figure 3C, Figures S3A-C). Since we primarily wanted to analyze the impact of tributyrin on gene expression, we focused our analysis on genes that changed with tributyrin treatment compared to ampicillin alone. We performed hierarchical clustering and observed four clusters with distinct patterns of gene expression (Figure 3C). While tributyrin impacted expression of some genes that were not affected by ampicillin (clusters 2 and 3), we reasoned that these genes may reflect a different mechanism than the potential action of butyrate on chromatin (i.e., binding of butyrate to cell surface receptors). We therefore focused on genes that displayed a pattern of expression that matched our observations of histone acyl marks: genes that were altered in expression with ampicillin treatment, and then rescued back to a baseline similar to the vehicle treated group (clusters 1 and 4, Figure 3C-D). In these clusters, tributyrin treatment resulted in significant enrichment in gene ontology categories related to metabolism. These observations are consistent with previously published roles of butyrate in energy metabolism (Donohoe, Wali, et al., 2012). Furthermore, we observed that tributyrin treatment resulted in the downregulation of genes related to mitochondria and oxidative phosphorylation, as well as autophagy, while upregulated genes were enriched in categories related to glutathione metabolism and catabolic processes (Figure 3E, Figure 3S). Together, these observations suggest that tributyrin results in increased histone butyrylation and gene expression changes that are enriched in gene programs related to metabolism.

To further investigate the landscape of histone acylations in IECs, we performed ChIP-seq using antibodies targeting different histone PTMs. Here, we again focused our analysis on the localization of H3K27bu. We observed that H3K27bu largely overlaps with other active histone modifications, including H3K27ac and H3K4me3 (Figure 4A-B). In contrast, H3K27me3, which is associated with gene repression, is localized to distinct regions. We also analyzed the genomic localization of H3K27bu peaks, and as expected from our visualization of ChIP-seq signal, this histone PTM is enriched at active gene regulatory elements (Figure 4C). Namely, H3K27bu peaks showed pronounced enrichment versus the genome at promoter regions, especially within 1 kb of the transcription start site (TSS), as 95.8% of H3K27bu peaks were localized to the immediate downstream promoter (compared to these sites comprising only 2.2% of genome). We next wanted to determine whether H3K27bu occupancy is related to overall levels of gene expression. In analyzing IECs from untreated mice, we binned our RNA-seq data into quartiles based on expression level. Interestingly, H3K27bu signal was highest in the quartile with highest gene expression and lowest in the quartile with lowest expression (Figure 4D). Thus, we conclude that H3K27bu is associated with active gene regulatory elements and gene expression in IECs.

**Figure 4:**
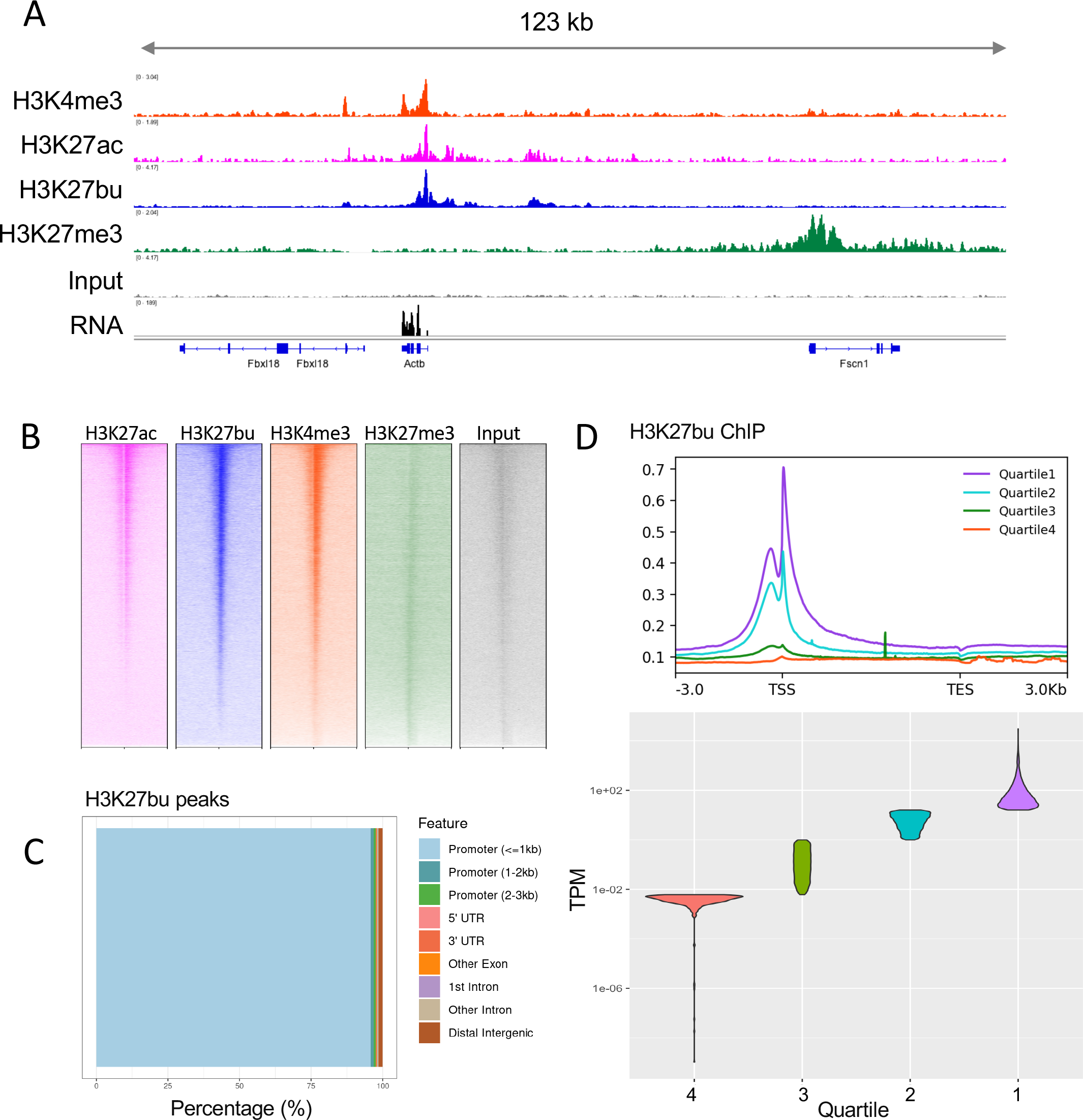
Histone butyrylation on H3K27 is associated with active gene regulatory elements and gene expression in intestinal epithelial cells. (A) Representative tracks of ChIP-seq data in intestinal epithelial cells. At least two biological replicates were performed for each ChIP. A representative RNA-seq track is also shown (n = 3). (B) Heatmaps of ChIP-seq signals centered around transcription start sites (TSSs). Range for each heatmap is −2kb to +2kb around the TSS. (C) Genomic localization of consensus H3K27bu peaks of two replicates by percentage. (D) H3K27bu localizes to highly expressed genes. Meta plots of a representative H3K27bu ChIP signal shown above violin plot displaying the quartiles of gene expression.

## DISCUSSION

Our studies have identified and shown the functional importance of select histone PTMs in a physiological context. Namely, we detected histone butyrylation and propionylation on histone H3 lysines 9 and 27. These non-acetyl histone acyl marks are localized in mammalian intestinal epithelial cells, along with more traditionally studied PTMs, including acetylation. Importantly, these PTMs are regulated by the microbiota, and in the case of histone butyrylation, the availability of the metabolite butyrate. This suggests that the landscape of histone PTMs is diverse in the intestine and can be regulated by the availability of donor molecules for these different PTMs. In addition, changes in histone butyrylation may be part of a mechanism of action of butyrate that has so far been overlooked.

The connection between how metabolism regulates histone PTMs and vice versa is an open question in the field. Some studies have suggested that histone PTMs can function as a reservoir of metabolites, where excess metabolites are deposited onto chromatin as histone PTMs for storage, and then recycled by chromatin modifying enzymes (Mendoza et al., 2022; Ye & Tu, 2018). This hypothesis would fit with our study, where both butyrate and acetate are in millimolar concentrations in the intestinal lumen and likely accumulate in cells as butyryl-CoA and acetyl-CoA, respectively. Investigating this hypothesis may be an important course of future study. However, the metabolic reservoir hypothesis is also not mutually exclusive from these PTMs also having gene regulatory functions.

A major question from our work is: what is the function of histone butyrylation in gene expression, specifically in the intestinal epithelium? Interestingly, our RNA-seq results suggested that the majority of genes rescued by tributyrin treatment in mice are downregulated (Figure 3C). This may be surprising, as many studies have found that histone acyl marks are activating in transcription, including butyrate on histone H4 (Goudarzi et al., 2016). In addition, we find that H3K27bu levels are correlated with gene expression (Figure 4D). However, a study from the Strahl and Morrison laboratories provides a contrasting view, in which they describe a repressive function of histone crotonylation during yeast metabolic cycling (Gowans et al., 2019). These paradoxical observations suggest that more studies are needed to determine the functional role of H3K27bu *in vivo*. In addition, using traditional RNA-seq gives us a snapshot of gene expression in time, yet we have not determined what may be direct versus indirect effects of H3K27bu. We speculate that investigating transcription factor-mediated responses and reader proteins (which are often effectors for histone PTMs) for these histone acylations and their potential impact on chromatin structure may be key future directions.

One limitation of our study is that we did not separate the effects of histone butyrylation vs. acetylation. Sodium butyrate is a well-characterized HDAC inhibitor, and we recognize that butyrate as an HDAC inhibitor has important biological roles, both in intestinal epithelial cells and other cell populations in the gut (Furusawa et al., 2013; Wu et al., 2020). Due to this activity in inhibiting HDACs, butyrate treatment induces a concurrent increase in both histone acetylation and butyrylation. The use of additional model systems would be especially useful to tease apart the effects of these two similar histone PTMs, such as using cell culture models in which chromatin writers selectively utilize butyryl-CoA vs. acetyl-CoA (Liu et al., 2017). Furthermore, our analyses are focused on gene expression in bulk epithelial cells from the intestine, which has a complex architecture and distinct heterogeneous cell types. Key next steps include determining how the IEC response to butyrate and other microbial metabolites varies between these different specialized cell types.

## ACKNOWLEDGEMENTS

We thank members of the Allis and Mucida laboratories, Kivanç Birsoy, Steven Josefowicz, and Rachel Niec, for helpful discussions and support. C.D.A. wishes to acknowledge the encouragement of the above Tri-Institutional faculty in our pursuing these studies and The Rockefeller University for financial support. We thank Alexey Soshnev for assistance with figure design and utilized BioRender for additional schematics. L.A.G. was supported by the Shapiro-Silverberg Fund for the Advancement of Translational Research at The Rockefeller University, a National Institutes of Health (NIH) Ruth L. Kirschstein NRSA fellowship (F32GM134560), and a NIH MOSAIC Career Award (K99GM143550). P.J.L. was supported by the Crohn’s and Colitis Foundation (RFA 598467 to P.J.L) and the NIH (T32CA009140). B.A.G was supported by the NIH (R01AI118891 and R01HD106051 to B.A.G.). We acknowledge the help and support of the following Resource Centers at Rockefeller University: Bio-Imaging, Genomics, Bioinformatics, and Comparative Bioscience Center.

## AUTHOR CONTRIBUTIONS

L.A.G. and C.D.A. conceptualized this project and wrote the manuscript with input from all authors. L.A.G., B.S.R., P.J.L., M.L., M.P., and Z.N. conducted experiments, contributed to sample preparation, and provided conceptual advice. T.S.C., B.A.G., D.M., and C.D.A. participated in study supervision.

## EXPERIMENTAL PROCEDURES

### Animals

C57BL/6J mice were purchased from Jackson Laboratories (000664). B6NTac murine pathogen free (MPF) or germ-free (GF) mice were purchased from Taconic Biosciences. All animals utilized were between 6-10 weeks old. Female mice were used for experiments unless otherwise noted. For treatment with antibiotics, single or multiple antibiotics (1 g/L ampicillin, 1 g/L neomycin, 0.5 g/L metronidazole, and/or 0.5 g/L vancomycin) were dissolved in drinking water with 10 g/L Splenda. Mice were acclimated to water with Splenda alone for two days prior to administration of antibiotics. Gavages were performed using either 200 ml tributyrin (Sigma) or with 200 ml equimolar glycerol solution for the mock gavage groups. Animal care and use followed NIH guidelines and was approved by the Institutional Animal Care and Use Committee (IACUC) at The Rockefeller University.

### Histone extractions from tissues

All tissue samples were collected at time of sacrifice and snap frozen in liquid nitrogen and stored at −80 °C until processing. All tissues of the gastrointestinal tract were first minced in cold PBS with protease inhibitors. The tissue was spun briefly and the pellet brought up in cold PBS with protease inhibitors and 0.1% NP-40. Tissue homogenization was carried out using a Pellet Pestle for approximately 20-30 seconds per sample. Samples were then centrifuged at 200×*g* for 5 minutes and the supernatant removed. The pellet was then brought up in Modified RIPA Buffer (50 mM Tris-HCL pH 8.0, 150 mM NaCl, 1 mM EDTA, 1% Triton-X, 0.25% Sodium Deoxycholate, 0.1% SDS, Complete Protease Inhibitor, and 1 mM PMSF) and incubated on ice for 20 minutes. Samples were then centrifuged for 10 minutes at 16,000×*g*. The supernatant was saved as tissue lysate, and the pellet was processed for acid extraction of histones (Shechter et al., 2007). Briefly, pellets were vortexed in 0.4 N H_2_SO_4_ and incubated overnight, then precipitated with 100% TCA. After precipitation, pellets were washed with acetone and allowed to air dry. Histones were then resuspended in water and protein concentration was assessed with BCA assay (Thermo) and Coomassie staining.

### Mass spectrometry

Histones were prepared for MS analysis using derivatization with propionic anhydride as previously described (Lund et al., 2019) except that isotopically labeled propionic anhydride (D10, 98%, Cambridge Isotope Laboratories Inc.) was used to differentiate endogenous (light) from artificial (heavy) histone propionylation. Briefly, 1 volume of 25% propionic anhydride in 2-propanol was mixed with 2 volumes of 0.1M ammonium bicarbonate containing 10-20 μg of histone extracts or 2 μg of mono-nucleosome extracts. After allowing the reaction to proceed for 15 mins at 37°C, the histones were dried in a speed-vac. This derivatization was performed once more prior to digestion with trypsin (1 μg per 20 μg histones) overnight at room temperature in 0.1M ammonium bicarbonate. After digestion, two additional rounds of derivatization were performed to propionylate the free N-termini. Peptides were then desalted with C18 stage tips for analysis by LC-MS/MS.

Peptides were resolved by reversed phase liquid chromatography using EASY-nLC 1000, Dionex Ultimate3000, or NeoVanquish (Thermo) systems fitted with 75 μm i.d. columns (either 15-20 cm packed in-house with 2.4 μm C18 material or 25 cm Acclaim PepMap 3 μm from Thermo) positioned in line with a QE or QE-HF mass spectrometer (Thermo). The chromatography gradient typically started at 2% B for 2 mins, increasing to 10% B from 2-4 mins, increasing to 30% B from 4-49 mins, and increasing to 90% B from 49-52 mins. In some cases, another gradient was used increasing from 2 to 45% B from 0-40 mins and then increasing to 95% B from 40-41 mins. Solvents A and B consisted of 0.1% formic acid in water and 80% acetonitrile with 0.1% formic acid, respectively. Parallel reaction monitoring (PRM) was used to analyze peptides of interest, including the unmodified, acetylated, butyrylated, and endogenously propionylated forms of H3K9 and H3K27. Typical parameters of the PRM were as follows: positive polarity; 17,500 or 30,000 resolution; 5e5 or 1e6 AGC; 50 or 100 ms max injection time; 11 or 20 loop count; 1.2 m/z isolation width or 3.0 m/z isolation width with 0.7 m/z offset; 30 NCE; centroid mode. Each cycle also included a full MS scan with typical settings as follows: positive polarity; 60,000 or 70,000 resolution; 1e6 AGC; 50 or 100 ms max injection time; 200-1200 or 250-1000 m/z scan range; profile mode. Data were analyzed with Skyline and fragmentation patterns were visualized with the Interactive Peptide Spectral Annotator (interactivepeptidespectralannotator.com, (Brademan et al., 2019)). Peptide modifications that were considered included artificial D5-propionyl (+61.057598 to K and N-termini), endogenous propionyl (+56.026215 to K), butyryl (+70.041865 to K), and acetyl (+42.010565 to K).

To construct the titration curve of histone acyl marks, 2 μg of unmodified or acylated mono-nucleosomes (H3K9ac, H3K9bu, H3K27ac, H3K27bu) were separately propionylated, digested, and analyzed by LC-MS/MS using 0.2 μg injections. Based on the observed peak areas of selected unmodified peptides from H2A, H2B, H3, and H4, peptide concentrations were equalized again to 110 ng/μl. A 10-fold dilution series was then prepared from a standard mixture of acyl marks, originally containing 3 μl of each acyl mark. 8 μl of each dilution, including the starting mixture, was then combined with 2 μl of peptides from unmodified nucleosomes. In this way, accounting for the unmodified H3K9 and H3K27 present in acylated H3K27 or H3K9 mono-nucleosomes, respectively, in addition to the unmodified H3K9 and H3K27 in unmodified mono-nucleosomes, the ratio of each acyl mark to its corresponding unmodified background ranges from 3.3E-1 to 2E-7. Final peptide concentrations were equalized to 22 ng/μl with solvent and 4.5 μl (100 ng) injections were used for LC-MS/MS analysis.

### Immunoblotting

Samples were run on 16% or 4-20% tris glycine SDS-PAGE gels (Invitrogen) and transferred to PVDF membranes. Blocking and all antibody incubations were done in 5% milk, with TBST washes in between. Immunoblots were imaged using Immobilon ECL (Millipore) and an Amersham Imager 600 (GE). Antibodies used are as follows: pan-acetyllysine (PTM-Biolabs PTM-105), pan-butyryllysine (PTM-Biolabs PTM-301), pan-propionyllysine (PTM-Biolabs PTM-201), H3 (Abcam, ab1791), H3K27bu (Millipore ABE2854), H3K9bu (PTM-Biolabs PTM-305), H3K27pr (Millipore ABE2853), H3K9pr (Millipore ABE2852), H3K27ac (Active Motif 39133).

### Immunofluorescence

Intestinal tissues were fixed with 2% paraformaldehyde in PBS overnight. After fixation, tissues were washed with B1n buffer, pH 7.0 (0.3 M Glycine, 0.1% v/v Triton-X 100) followed by PBS, and then stored in 70% ethanol. Tissues were embedded in parafilm and sectioned to 5 μm sections by HistoWiz, Inc. After rehydrating, sections were boiled in sodium citrate buffer, pH 6.0 (10 mM sodium citrate, 0.05% Tween 20), and then stained. Imaging was performed on a Zeiss Inverted LSM 780 laser scanning confocal microscope using a 40x objective. Stainings and dilutions: α-H3K27bu (Millipore ABE2854, 1:500), α-Villin-647 (SantaCruz sc58897-647, 1:200), Goat anti-Rabbit Alexa Fluor 488 (1:1,000), DAPI (1:1,000).

### Intestinal epithelial cell isolation

Epithelial cell isolation was carried out as previously described and further optimized for the cecum (Gracz et al., 2012; Reis et al., 2013). All steps were performed on ice or 4 °C unless otherwise noted. After excising cecum and/or colon tissue, any visible fat was removed. Tissues were then opened longitudinally and cleaned of feces with sequential washing in PBS. Once all visible feces were removed and tissues appeared clean, tissues were cut into approximately 1 cm sections and placed in Dissociation Reagent 1 (30 mM EDTA, 1.5 mM DTT in PBS) on ice for 20 minutes. Tissues were then transferred to Dissociation Reagent 2 (30 mM EDTA in DMEM + 2% FCS) and incubated at 37 °C for 10 minutes. After incubation, tissues were shaken by hand for 30 seconds until the solution became cloudy. The supernatant was then filtered through a stainless-steel sieve and spun at 600×*g* for 5 mins. The pellet was then washed with PBS + 10% FBS and centrifuged again. Finally, the pellet was resuspended in 10 mL HBSS with 0.5 U/ml Dispase (Corning) and incubated for 5-10 minutes at 37 °C, shaking every 2 mins. Cells were checked for dissociation starting at 5 minutes, during which samples were placed back on ice. Once mostly single cells were observed, 10% FBS was added to quench the reaction. Cells were sequentially pass cells through 70 μm and 40 μm filters, spun down and washed once with 10% FBS. The final pellet was resuspended in FACS buffer (2.5% FBS in PBS), and cells were counted and visualized using a hemocytometer.

### Chromatin Immunoprecipitation

Isolated IECs were fixed in PBS with 1% formaldehyde for 10 minutes, followed by quenching with 125 mM glycine for 5 minutes. Crosslinked cells were washed once with PBS and then lysed using the NEXSON protocol (Arrigoni et al., 2016). Briefly, cells were resuspended in FL Buffer (5 mM PIPES, pH 8.0, 85 mM KCl, 0.5% NP-40, protease inhibitor cocktail) and sonicated in the Bioruptor Pico (Diagenode) on low power at 15 seconds on 30 seconds off cycles until at least 70% cell membranes lysed and intact nuclei were visible. Nuclei were then washed in FL Buffer and resuspended in D3 Sonication Buffer (10 mM Tris-HCl, pH 8.0, 0.1% SDS, 1 mM EDTA, protease inhibitor cocktail) for sonication using the Covaris E220 (Covaris). Sonication was optimized in an assay-dependent manner and input DNA was checked prior to proceeding with immunoprecipitations (IPs). For IPs, sonicated chromatin was diluted into ChIP Dilution Buffer (10 mM Tris-HCl, pH 8.0, 1 mM EDTA, 150 mM NaCl, 1% Triton-X, protease inhibitor cocktail) and added to antibody-bound Protein A Dynabeads (Invitrogen) and incubated overnight in the cold room, rotating. Antibodies used were as follows: H3K27ac (Active Motif 39133), H3K27bu (Millipore ABE2854), H3K4me3 (Active Motif 39159), H3K27me3 (Cell Signaling 9733). The next day, beads were washed six times with RIPA buffer (50 mM HEPES-KOH, pH 7.5, 100 mM LiCl, 1 mM EDTA, 0.7% Na-Deoxycholate, 1% NP-40) followed by one wash with TE-NaCl Buffer (10 mM Tris-HCl, pH 8.0, 50 mM NaCl, 1 mM EDTA). DNA was eluted from the beads using Elution Buffer (50 mM Tris-HCl, pH 8.0, 10 mM EDTA, 1% SDS). DNA was reverse crosslinked at 65 °C overnight, followed by RNase and ProteinaseK digestion and purification using Phenol-Chloroform Isoamyl Alcohol mix (Millipore) and PhaseLock tubes (5PRIME). DNA was quantified using Qubit 4 Fluorometer (Thermo) prior to proceeding with library preparation.

### RNA isolation

Total RNA was isolated from IECs using RNeasy Mini Kit (Qiagen) with on-column DNA digestion. RNA samples were analyzed on the Bioanalyzer RNA 6000 Pico (Agilent) prior to library preparation.

### Library preparations & sequencing

RNA libraries were converted to cDNA libraries using NEBNext Poly(A) mRNA Magnetic Isolation Module (NEB) and NEBNext Ultra II RNA Library Prep Kit for Illumina (NEB) according to the manufacturer’s instructions. ChIP libraries were prepared using NEBNext Ultra II DNA Library Prep Kit for Illumina (NEB) according to the manufacturer’s instructions. All libraries were analyzed by TapeStation prior to sequencing. Single-end sequencing was performed on the Illumina NextSeq 500 sequencer.

### Bioinformatics analysis

Sequence and transcript coordinates for mouse mm10 UCSC genome and gene models were retrieved from the Bioconductor Bsgenome.Mmusculus.UCSC.mm10 (version 1.4.0) and TxDb.Mmusculus.UCSC.mm10.knownGene (version 3.4.0) Bioconductor libraries respectively.

For the analysis of RNAseq data, transcript expressions were calculated using the Salmon quantification software (Patro et al., 2017)(version 0.8.2) and gene expression levels as TPMs and counts retrieved using Tximport (Love et al., 2016)(version 1.8.0). Normalization and rlog transformation of raw read counts in genes, PCA and differential gene expression analysis were performed using DESeq2 (Love et al., 2016)(version 1.20.0). Additional sample to sample variability assessment was made with heat maps of between sample distances using Pheatmap (version 1.0.10). Significant genes (padj < 0.05) from a pairwise comparison between mice treated with ampicillin plus tributyrin vs. ampicillin with mock gavage were used for hierarchical clustering. The Z-score of the rlog of gene counts was used as the input for clustering. Clustering was done using Pheatmap after determining optimal number of clusters using the elbow method. Gene set enrichment tests were conducted using clusterProfiler (version 3.18.1) (Yu et al., 2012).

For the analysis of ChIPseq data, reads were mapped using the Rsubread package’s align function (version 1.30.6) (Liao et al., 2019). Peaks of enrichment were determined using the MACS peak caller (version 2.1.1) (Y. Zhang et al., 2008). Consensus peaks were determined to be peaks that are found in common across replicates. Peaks were annotated and genome distribution was determined using the ChIPseeker package (Version 1.26.2) (Yu et al., 2015). Heatmaps and metaplots were generated with deeptools (version 3.5). Normalized, fragment extended signal bigWigs are created using the rtracklayer package (version 1.40.6), and then visualized and exported from IGV.

### Statistics

Details for statistical tests and replicates are described in the figure legends. Prism 7 (GraphPad) was used to generate plots and perform statistical tests. Error bars represent the standard error. Unpaired two-tailed Student’s t-test or one-way ANOVA with multiple comparisons was used to assess significance and is indicated in the figure legends. P<0.05 was considered statistically significant.

## Supplemental Information

**Figure S1:**
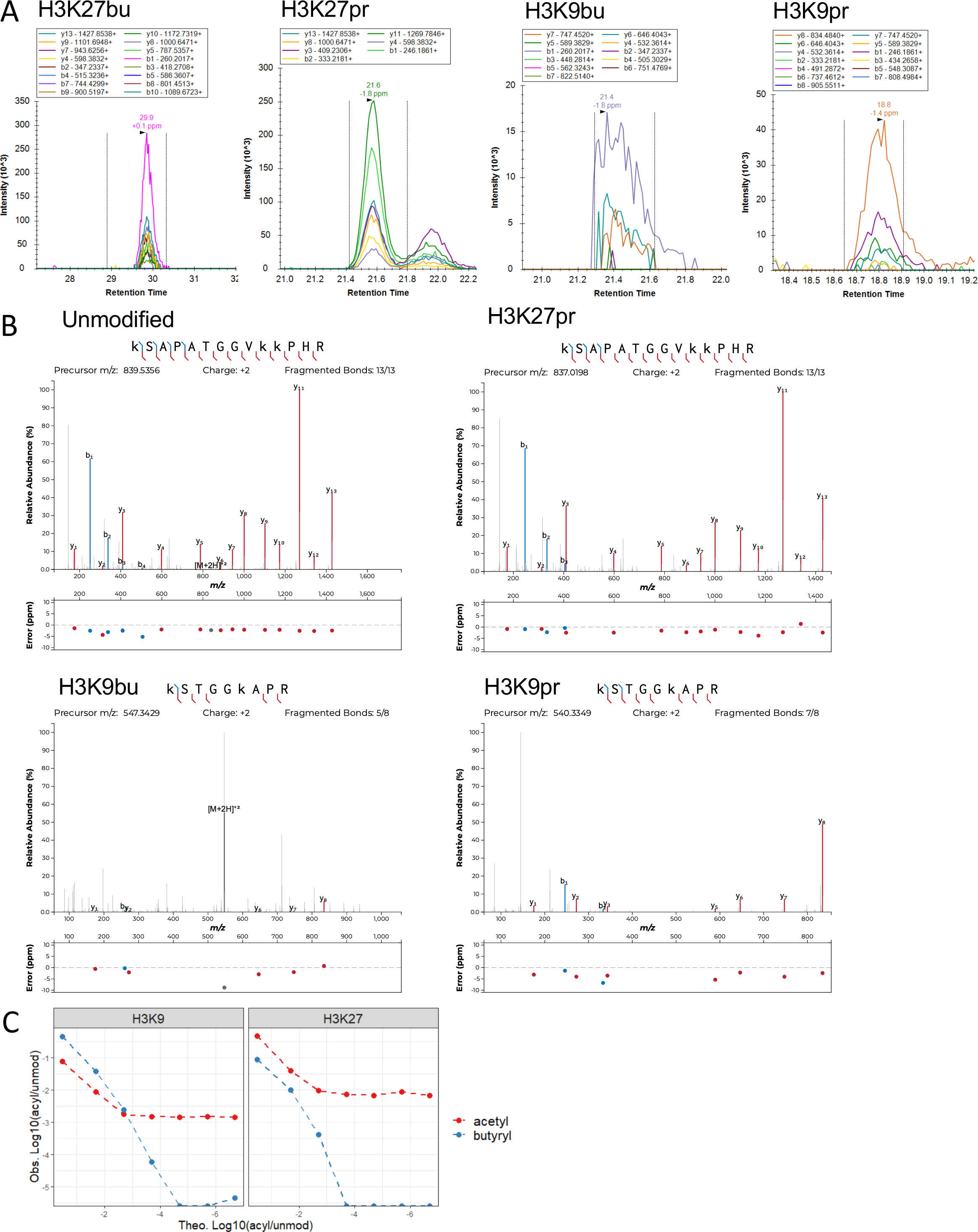
Identification of histone acylations in the mouse intestine. (A) Chromatograms of fragment ions from precursor histone peptides with acyl marks from mouse cecum. Representative traces are shown out of n = 3 mice. (B) Fragmentation spectra of unmodified peptides and peptides with H3K27pr, H3K9bu, and H3K9pr. All representative spectra are from histones extracted from mouse cecal samples. (C) Butyrylated peptides are not as readily detected as acetylated peptides at low relative abundances as assessed by concomitant serial dilution of acetylated and butyrylated H3K9 and H3K27 nucleosomes into unmodified nucleosomes.

**Figure S2:**
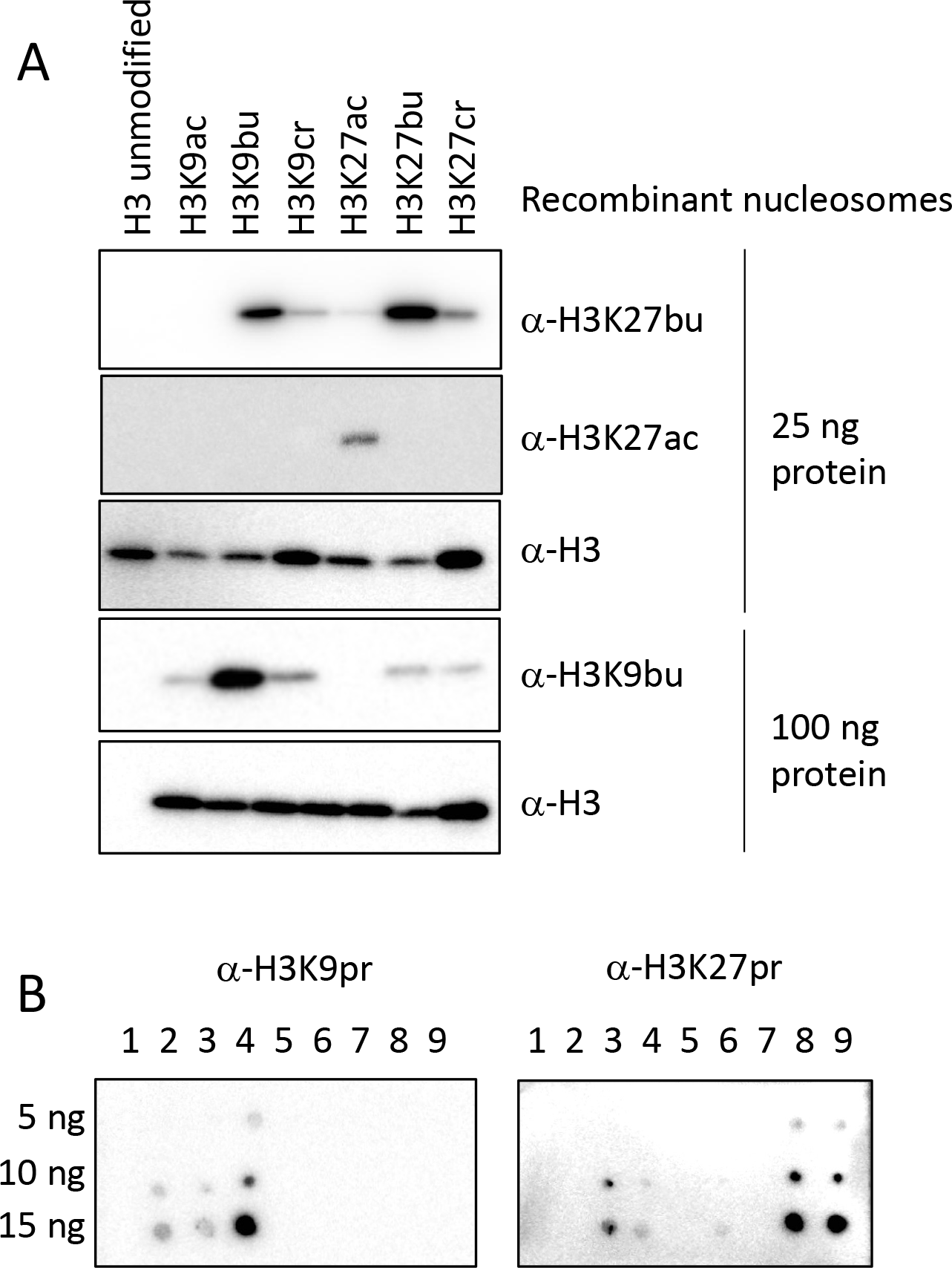
Characterization of acyl antibodies. Abbreviations are as follows: ac = acetyl, bu = butyryl, cr = crotonyl, and pr = propionyl. (A) Testing of antibody specificity using recombinant nucleosomes. 30 or 100 ng recombinant nucleosomes were run on an SDS-PAGE gel and subjected to immunoblotting. (B) Testing of propionyl antibody specificity by dot blotting. 5, 10, or 15 ng of H3 peptides were placed on nitrocellulose membrane and then incubated with the indicated antibodies. Lanes: 1) H3 1-20 unmodified, 2) H3K9ac, 3) H3K9bu, 4) H3K9pr, 5) H3.1 15-34 unmodified, 6) H3.1 K27ac, 7) H3.3 15-34 unmodified, 8) H3.3 K27bu, 9) H3.3 K27pr.

**Figure S3:**
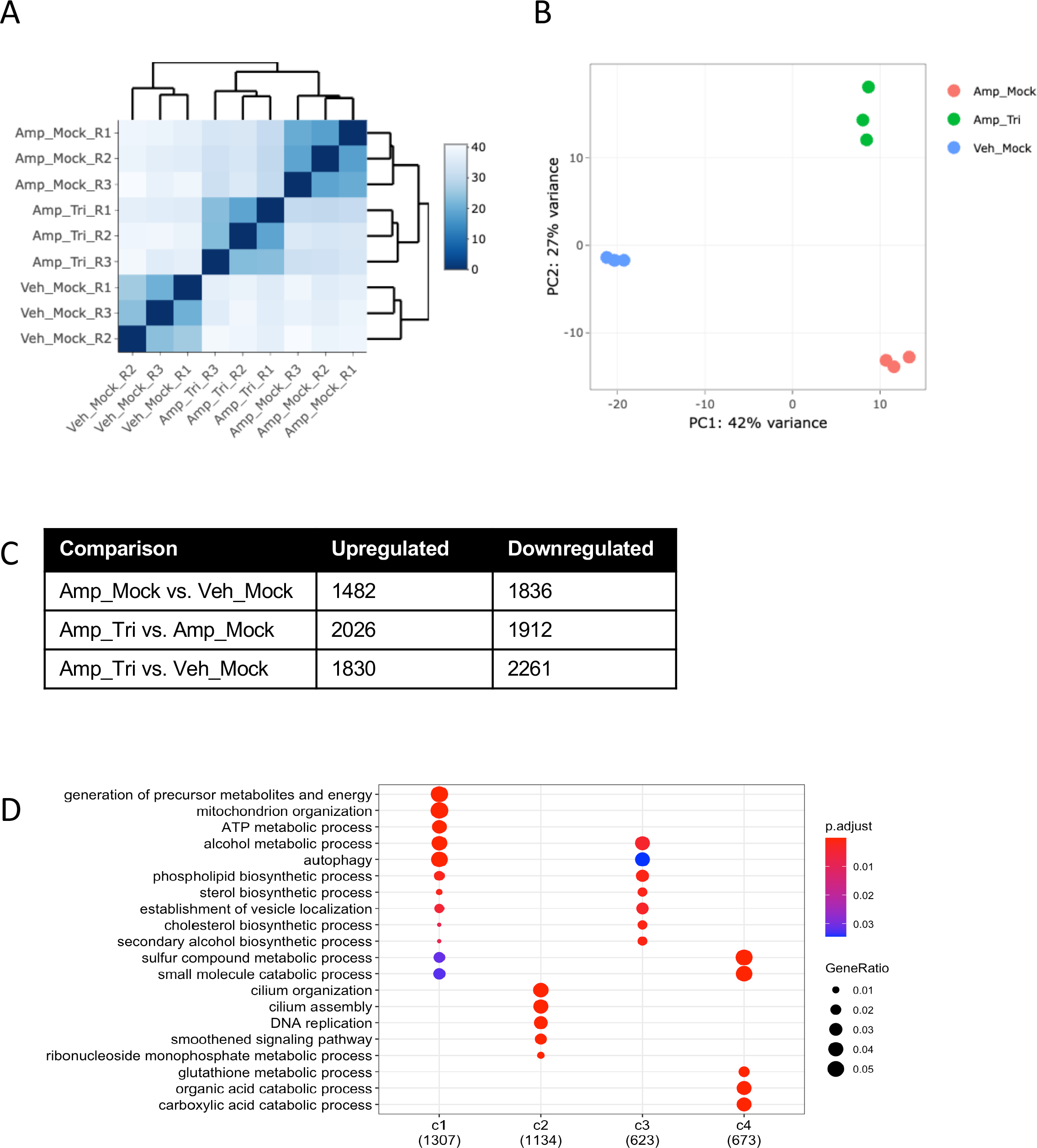
Characterization of RNA-seq data. Mice were divided into three different groups (n= 3 per group): vehicle treated and mock gavaged (Veh_Mock), Ampicillin treated and mock gavaged (Amp_Mock), and Ampicillin treated with tributyrin gavage (Amp_Tri). IECs were isolated for RNA sequencing. (A) Heatmap of correlation between samples. Pearson correlation was performed of gene expression measurements across samples using DEseq2. (B) Principal component analysis of RNA-seq samples. (C) Table of significant differential gene expression across groups. Gene expression changes were identified using DEseq2 and p-values less than 0.05 were considered significant. (D) Gene ontology analysis of all clusters of differential gene expression.

## REFERENCES

Allis, C. D. & Jenuwein, T. (2016). The molecular hallmarks of epigenetic control. Nature Reviews Genetics, 17(8), 487–500. https://doi.org/10.1038/nrg.2016.59

Arrigoni, L., Richter, A. S., Betancourt, E., Bruder, K., Diehl, S., Manke, T. & Bönisch, U. (2016). Standardizing chromatin research: a simple and universal method for ChIP-seq. Nucleic Acids Research, 44(7), e67–e67. https://doi.org/10.1093/nar/gkv1495

Brademan, D. R., Riley, N. M., Kwiecien, N. W. & Coon, J. J. (2019). Interactive Peptide Spectral Annotator: A Versatile Web-based Tool for Proteomic Applications. Molecular & Cellular Proteomics, 18(8), S193–S201. https://doi.org/10.1074/mcp.tir118.001209

Brown, A. J., Goldsworthy, S. M., Barnes, A. A., Eilert, M. M., Tcheang, L., Daniels, D., Muir, A. I., Wigglesworth, M. J., Kinghorn, I., Fraser, N. J., Pike, N. B., Strum, J. C., Steplewski, K. M., Murdock, P. R., Holder, J. C., Marshall, F. H., Szekeres, P. G., Wilson, S., Ignar, D. M., … Dowell, S. J. (2003). The Orphan G Protein-coupled Receptors GPR41 and GPR43 Are Activated by Propionate and Other Short Chain Carboxylic Acids. Journal of Biological Chemistry, 278(13), 11312–11319. https://doi.org/10.1074/jbc.m211609200

Brownell, J. E., Zhou, J., Ranalli, T., Kobayashi, R., Edmondson, D. G., Roth, S. Y. & Allis, C. D. (1997). Tetrahymena Histone Acetyltransferase A: A Homolog to Yeast Gcn5p Linking Histone Acetylation to Gene Activation. Cell, 84, 843–851. https://doi.org/10.1016/s0092-8674(00)81063-6

Carey, B. W., Finley, L. W. S., Cross, J. R., Allis, C. D. & Thompson, C. B. (2015). Intracellular α-ketoglutarate maintains the pluripotency of embryonic stem cells. Nature, 518(7539), 413–416. https://doi.org/10.1038/nature13981

Cummings, J. H., Pomare, E. W., Branch, W. J., Naylor, C. P. & Macfarlane, G. T. (1987). Short chain fatty acids in human large intestine, portal, hepatic and venous blood. Gut, 28(10), 1221. https://doi.org/10.1136/gut.28.10.1221

Dai, Z., Ramesh, V. & Locasale, J. W. (2020). The evolving metabolic landscape of chromatin biology and epigenetics. Nature Reviews Genetics. https://doi.org/10.1038/s41576-020-0270-8

den Besten, G. den, Eunen, K. van, Groen, A. K., Venema, K., Reijngoud, D.-J. & Bakker, B. M. (2013). The role of short-chain fatty acids in the interplay between diet, gut microbiota, and host energy metabolism. Journal of Lipid Research, 54(9), 2325–2340. https://doi.org/10.1194/jlr.r036012

Donohoe, D. R., Collins, L. B., Wali, A., Bigler, R., Sun, W. & Bultman, S. J. (2012). The Warburg Effect Dictates the Mechanism of Butyrate-Mediated Histone Acetylation and Cell Proliferation. Molecular Cell, 48(4), 612–626. https://doi.org/10.1016/j.molcel.2012.08.033

Donohoe, D. R., Wali, A., Brylawski, B. P. & Bultman, S. J. (2012). Microbial Regulation of Glucose Metabolism and Cell-Cycle Progression in Mammalian Colonocytes. PLoS ONE, 7(9), e46589–9. https://doi.org/10.1371/journal.pone.0046589

Farrelly, L. A., Thompson, R. E., Zhao, S., Lepack, A. E., Lyu, Y., Bhanu, N. V., Zhang, B., Loh, Y.-H. E., Ramakrishnan, A., Vadodaria, K. C., Heard, K. J., Erikson, G., Nakadai, T., Bastle, R. M., Lukasak, B. J., Zebroski, H., Alenina, N., Bader, M., Berton, O., … Maze, I. (2019). Histone serotonylation is a permissive modification that enhances TFIID binding to H3K4me3. Nature, 567(7749), 535–539. https://doi.org/10.1038/s41586-019-1024-7

Fellows, R., Denizot, J., Stellato, C., Cuomo, A., Jain, P., Stoyanova, E., Balázsi, S., Hajnády, Z., Liebert, A., Kazakevych, J., Blackburn, H., Corrêa, R. O., Fachi, J. L., Sato, F. T., Ribeiro, W. R., Ferreira, C. M., Perée, H., Spagnuolo, M., Mattiuz, R., … Varga-Weisz, P. (2018). Microbiota derived short chain fatty acids promote histone crotonylation in the colon through histone deacetylases. Nature Communications, 1–15. https://doi.org/10.1038/s41467-017-02651-5

Fleming, S. E., Fitch, M. D., DeVries, S., Liu, M. L. & Kight, C. (1991). Nutrient Utilization by Cells Isolated from Rat Jejunum, Cecum and Colon. The Journal of Nutrition, 121(6), 869–878. https://doi.org/10.1093/jn/121.6.869

Furusawa, Y., Obata, Y., Fukuda, S., Endo, T. A., Nakato, G., Takahashi, D., Nakanishi, Y., Uetake, C., Kato, K., Kato, T., Takahashi, M., Fukuda, N. N., Murakami, S., Miyauchi, E., Hino, S., Atarashi, K., Onawa, S., Fujimura, Y., Lockett, T., … Ohno, H. (2013). Commensal microbe-derived butyrate induces the differentiation of colonic regulatory T cells. Nature, 504(7480), 446–450. https://doi.org/10.1038/nature12721

Goudarzi, A., Zhang, D., Huang, H., Barral, S., Kwon, O. K., Qi, S., Tang, Z., Buchou, T., Vitte, A.-L., He, T., Cheng, Z., Montellier, E., Gaucher, J., Curtet, S., Debernardi, A., Charbonnier, G., Puthier, D., Petosa, C., Panne, D., … Khochbin, S. (2016). Dynamic Competing Histone H4 K5K8 Acetylation and Butyrylation Are Hallmarks of Highly Active Gene Promoters. Molecular Cell, 62(2), 169–180. https://doi.org/10.1016/j.molcel.2016.03.014

Gowans, G. J., Bridgers, J. B., Zhang, J., Dronamraju, R., Burnetti, A., King, D. A., Thiengmany, A. V., Shinsky, S. A., Bhanu, N. V., Garcia, B. A., Buchler, N. E., Strahl, B. D. & Morrison, A. J. (2019). Recognition of Histone Crotonylation by Taf14 Links Metabolic State to Gene Expression. Molecular Cell, 76(6), 909–921.e3. https://doi.org/10.1016/j.molcel.2019.09.029

Gracz, A. D., Puthoff, B. J. & Magness, S. T. (2012). Somatic Stem Cells, Methods and Protocols. Methods in Molecular Biology, 879, 89–107. https://doi.org/10.1007/978-1-61779-815-3_6

Kebede, A. F., Nieborak, A., Shahidian, L. Z., Gras, S. L., Richter, F., Gómez, D. A., Baltissen, M. P., Meszaros, G., Magliarelli, H. de F., Taudt, A., Margueron, R., Colomé-Tatché, M., Ricci, R., Daujat, S., Vermeulen, M., Mittler, G. & Schneider, R. (2017). Histone propionylation is a mark of active chromatin. Nature Structural & Molecular Biology, 24(12), 1048–1056. https://doi.org/10.1038/nsmb.3490

Kelly, C. J., Zheng, L., Campbell, E. L., Saeedi, B., Scholz, C. C., Bayless, A. J., Wilson, K. E., Glover, L. E., Kominsky, D. J., Magnuson, A., Weir, T. L., Ehrentraut, S. F., Pickel, C., Kuhn, K. A., Lanis, J. M., Nguyen, V., Taylor, C. T. & Colgan, S. P. (2015). Crosstalk between Microbiota-Derived Short-Chain Fatty Acids and Intestinal Epithelial HIF Augments Tissue Barrier Function. Cell Host & Microbe, 17(5), 662–671. https://doi.org/10.1016/j.chom.2015.03.005

Koh, A., Vadder, F. D., Kovatcheva-Datchary, P. & Bäckhed, F. (2016). From Dietary Fiber to Host Physiology: Short-Chain Fatty Acids as Key Bacterial Metabolites. Cell, 165(6), 1332–1345. https://doi.org/10.1016/j.cell.2016.05.041

Krautkramer, K. A., Kreznar, J. H., Romano, K. A., Vivas, E. I., Barrett-Wilt, G. A., Rabaglia, M. E., Keller, M. P., Attie, A. D., Rey, F. E. & Denu, J. M. (2016). Diet-Microbiota Interactions Mediate Global Epigenetic Programming in Multiple Host Tissues. Molecular Cell, 64(5), 982–992. https://doi.org/10.1016/j.molcel.2016.10.025

Liao, Y., Smyth, G. K. & Shi, W. (2019). The R package Rsubread is easier, faster, cheaper and better for alignment and quantification of RNA sequencing reads. Nucleic Acids Research, 47(8), e47–e47. https://doi.org/10.1093/nar/gkz114

Love, M. I., Hogenesch, J. B. & Irizarry, R. A. (2016). Modeling of RNA-seq fragment sequence bias reduces systematic errors in transcript abundance estimation. Nature Biotechnology, 34(12), 1287–1291. https://doi.org/10.1038/nbt.3682

Lund, P. J., Gates, L. A., Leboeuf, M., Smith, S. A., Chau, L., Friedman, E. S., Lopes, M., Saiman, Y., Kim, M. S., Petucci, C., Allis, C. D., Wu, G. D. & Garcia, B. A. (2021). Stable Isotope Tracing in vivo Reveals A Metabolic Bridge Linking the Microbiota to Host Histone Acetylation. BioRxiv, 2021.07.05.450926. https://doi.org/10.1101/2021.07.05.450926

Lund, P. J., Kori, Y., Zhao, X., Sidoli, S., Yuan, Z.-F. & Garcia, B. A. (2019). Isotopic Labeling and Quantitative Proteomics of Acetylation on Histones and Beyond (pp. 43–70). Springer New York. https://doi.org/10.1007/978-1-4939-9232-4_5

Mendoza, M., Egervari, G., Sidoli, S., Donahue, G., Alexander, D. C., Sen, P., Garcia, B. A. & Berger, S. L. (2022). Enzymatic transfer of acetate on histones from lysine reservoir sites to lysine activating sites. Science Advances, 8(3), eabj5688. https://doi.org/10.1126/sciadv.abj5688

Patro, R., Duggal, G., Love, M. I., Irizarry, R. A. & Kingsford, C. (2017). Salmon provides fast and bias-aware quantification of transcript expression. Nature Methods, 14(4), 417–419. https://doi.org/10.1038/nmeth.4197

Poul, E. L., Loison, C., Struyf, S., Springael, J.-Y., Lannoy, V., Decobecq, M.-E., Brezillon, S., Dupriez, V., Vassart, G., Damme, J. V., Parmentier, M. & Detheux, M. (2003). Functional Characterization of Human Receptors for Short Chain Fatty Acids and Their Role in Polymorphonuclear Cell Activation. Journal of Biological Chemistry, 278(28), 25481–25489. https://doi.org/10.1074/jbc.m301403200

Reis, B. S., Rogoz, A., Costa-Pinto, F. A., Taniuchi, I. & Mucida, D. (2013). Mutual expression of the transcription factors Runx3 and ThPOK regulates intestinal CD4+ T cell immunity. Nature Immunology, 14(3), 271–280. https://doi.org/10.1038/ni.2518

Roediger, W. E. W. (1982). Utilization of Nutrients by Isolated Epithelial Cells of the Rat Colon. Gastroenterology, 83(2), 424–429. https://doi.org/10.1016/s0016-5085(82)80339-9

Sabari, B. R., Tang, Z., Huang, H., Yong-Gonzalez, V., Molina, H., Kong, H. E., Dai, L., Shimada, M., Cross, J. R., Zhao, Y., Roeder, R. G. & Allis, C. D. (2015). Intracellular Crotonyl-CoA Stimulates Transcription through p300-Catalyzed Histone Crotonylation. Molecular Cell, 58, 203–215. https://doi.org/10.1016/j.molcel.2015.02.029

Sabari, B. R., Zhang, D., Allis, C. D. & Zhao, Y. (2016). Metabolic regulation of gene expression through histone acylations. Nature Reviews Molecular Cell Biology, 18, 90–101. https://doi.org/10.1038/nrm.2016.140

Shechter, D., Dormann, H. L., Allis, C. D. & Hake, S. B. (2007). Extraction, purification and analysis of histones. Nature Protocols, 2(6), 1445–1457. https://doi.org/10.1038/nprot.2007.202

Tan, M., Luo, H., Lee, S., Jin, F., Yang, J. S., Montellier, E., Buchou, T., Cheng, Z., Rousseaux, S., Rajagopal, N., Lu, Z., Ye, Z., Zhu, Q., Wysocka, J., Ye, Y., Khochbin, S., Ren, B. & Zhao, Y. (2011). Identification of 67 Histone Marks and Histone Lysine Crotonylation as a New Type of Histone Modification. Cell, 146(6), 1016–1028. https://doi.org/10.1016/j.cell.2011.08.008

Tognini, P., Murakami, M., Liu, Y., Eckel-Mahan, K. L., Newman, J. C., Verdin, E., Baldi, P. & Sassone-Corsi, P. (2017). Distinct Circadian Signatures in Liver and Gut Clocks Revealed by Ketogenic Diet. Cell Metabolism, 26(3), 523–538.e5. https://doi.org/10.1016/j.cmet.2017.08.015

Vidali, G., Boffa, L. C., Bradbury, E. M. & Allfrey, V. G. (1978). Butyrate suppression of histone deacetylation leads to accumulation of multiacetylated forms of histones H3 and H4 and increased DNase I sensitivity of the associated DNA sequences. Proceedings of the National Academy of Sciences, 75(5), 2239–2243. https://doi.org/10.1073/pnas.75.5.2239

Wellen, K. E., Hatzivassiliou, G., Sachdeva, U. M., Bui, T. V., Cross, J. R. & Thompson, C. B. (2009). ATP-Citrate Lyase Links Cellular Metabolism to Histone Acetylation. Science, 324(5930), 1076–1080. https://doi.org/10.1126/science.1167053

Wu, S., Hashimoto-Hill, S., Woo, V., Eshleman, E. M., Whitt, J., Engleman, L., Karns, R., Denson, L. A., Haslam, D. B. & Alenghat, T. (2020). Microbiota-derived metabolite promotes HDAC3 activity in the gut. Nature. https://doi.org/10.1038/s41586-020-2604-2

Ye, C. & Tu, B. P. (2018). Sink into the Epigenome: Histones as Repositories That Influence Cellular Metabolism. Trends in Endocrinology and Metabolism, 29(9), 626–637. https://doi.org/10.1016/j.tem.2018.06.002

Yu, G., Wang, L.-G., Han, Y. & He, Q.-Y. (2012). clusterProfiler: an R Package for Comparing Biological Themes Among Gene Clusters. OMICS: A Journal of Integrative Biology, 16(5), 284–287. https://doi.org/10.1089/omi.2011.0118

Yu, G., Wang, L.-G. & He, Q.-Y. (2015). ChIPseeker: an R/Bioconductor package for ChIP peak annotation, comparison and visualization. Bioinformatics, 31(14), 2382–2383. https://doi.org/10.1093/bioinformatics/btv145

Zhang, D., Tang, Z., Huang, H., Zhou, G., Cui, C., Weng, Y., Liu, W., Kim, S., Lee, S., Perez-Neut, M., Ding, J., Czyz, D., Hu, R., Ye, Z., He, M., Zheng, Y. G., Shuman, H. A., Dai, L., Ren, B., … Zhao, Y. (2019). Metabolic regulation of gene expression by histone lactylation. Nature, 574(7779), 575–580. https://doi.org/10.1038/s41586-019-1678-1

Zhang, Y., Liu, T., Meyer, C. A., Eeckhoute, J., Johnson, D. S., Bernstein, B. E., Nusbaum, C., Myers, R. M., Brown, M., Li, W. & Liu, X. S. (2008). Model-based Analysis of ChIP-Seq (MACS). Genome Biology, 9(9), R137. https://doi.org/10.1186/gb-2008-9-9-r137

